# DeepAge: Harnessing Deep Neural Network for Epigenetic Age Estimation From DNA Methylation Data of human blood samples

**DOI:** 10.1101/2024.08.12.607687

**Authors:** Sajib Acharjee Dip, Da Ma, Liqing Zhang

## Abstract

Accurate prediction of biological age from DNA methylation data is a critical endeavor in understanding the molecular mechanisms of aging and developing age-related disease interventions. Traditional epigenetic clocks rely on linear regression or basic machine learning models, which often fail to capture the complex, non-linear interactions within methylation data. This study introduces DeepAge, a novel deep learning framework utilizing Temporal Convolutional Networks (TCNs) to enhance the prediction of biological age from DNA methylation profiles using selected CpGs by a Dual-Correlation based apparoach. DeepAge leverages a sequence-based approach with dilated convolutions to effectively capture long-range dependencies between CpG sites, addressing the limitations of prior models by incorporating advanced network architectures including residual connections and dropout regularization. The dual correlation feature selection enhances our model’s predictive capabilities by identifying the most age-relevant CpG sites. Our model outperforms existing epigenetic clocks across multiple datasets, offering significant improvements in accuracy and providing deeper insights into the epigenetic determinants of aging. The proposed method not only sets a new standard in age estimation but also highlights the potential of deep learning in biologically relevant feature extraction and interpretation, contributing to the broader field of computational biology and precision medicine.

## Introduction

In the realm of biomedical research, the accurate estimation of biological age from epigenetic data, specifically DNA methylation, represents a pivotal challenge and opportunity. Biological age, as opposed to chronological age, offers a more nuanced view of an individual’s health status and aging process, informed by the epigenetic modifications that accumulate over time. These modifications, particularly methylation of DNA at CpG sites, have been robustly associated with various age-related changes and conditions. Traditional methods for estimating epigenetic age leverage linear and basic machine learning models, which, while foundational, often struggle with the complex, non-linear relationships inherent in methylation data across diverse biological systems. Various models such as Hannum (Hannum et al. 2013), Horvath1 (Horvath 2013), Horvath2 (Horvath et al. 2018), Lin (Lin et al. 2016), PhenoAge (Levine et al. 2018), and others employ diverse computational strategies, predominantly linear regression-based approaches that focus on weighted sums of methylation levels at selected CpG sites. These traditional models, while foundational, often do not account for the complex, non-linear interactions between CpGs that might influence aging processes more profoundly. For instance, Horvath’s clocks use linear algorithms that may not capture the entire spectrum of biological aging changes, leading to limitations in prediction accuracy across diverse populations and tissues. There are some non-linear parametric regression based methods for example, GP-Age (Varshavsky et al. 2023) which improve over flexible prediction but still suffer from capturing complex interaction.

Recent advancements have seen the application of more sophisticated machine learning techniques, such as random forests (Breiman 2001) and gradient boosting machines (Chen and Guestrin 2016), which have provided incremental improvements and more flexibility over linear models. However, these methods still often fall short of capturing the deeper interactions within methylation profiles. There are some methods applying deep learning appoaches, for example, PerSEClock (Zhao et al. 2024) applied channel attention, CPFNN (Li et al. 2021) used correlation pre-filtered neural network, MSCAP (Wang et al. 2023) used multi-scale CNN model, but most of them lack in considering their sequential or collective influence on aging that can be critical for a precise age prediction. Furthermore, most existing models have not fully explored the potential of deep learning techniques, which have revolutionized fields such as image and speech recognition for detecting intricate patterns in high-dimensional data.

Our work introduces DeepAge, a novel deep learning framework specifically designed to address these challenges in the context of epigenetic age estimation. DeepAge utilizes Temporal Convolutional Networks (TCNs) (Lea et al. 2016), which are particularly adept at handling sequence data, to model the sequential nature of CpG sites across the genome. This approach allows for an effective capture of long-range dependencies and interactions between CpG sites, which are essential for understanding the complex biological processes underlying aging. By integrating layers of temporal blocks that include dilated convolutions (Yu and Koltun 2015), DeepAge can access a broader context of the input sequence, thus enhancing its ability to discern pertinent aging signals from the methylation patterns. We implemented a dual correlation technique, utilizing both Spearman and Pearson correlations to identify CpGs most associated with age (De Winter, Gosling, and Potter 2016). By focusing on these relevant features, our model’s performance improved significantly, reducing the risk of overfitting and enhancing generalizability.

Moreover, DeepAge incorporates advanced techniques such as residual connections (He et al. 2016) and dropout regularization (Srivastava et al. 2014) to refine its learning process and avoid overfitting, a common challenge in deep learning models dealing with high-dimensional biological data. The architecture is designed to progressively increase its receptive field without inflating the model size excessively, making it both powerful and computationally efficient. This allows DeepAge not only to outperform existing epigenetic clocks (Hannum et al. 2013; Horvath 2013; Horvath et al. 2018; Lin et al. 2016; Belsky et al. 2022) in terms of prediction accuracy but also to provide deeper insights into the epigenetic factors that drive biological aging.

In summary, DeepAge represents a significant step forward in the field of epigenetics, offering a robust, scalable, and interpretable tool for age estimation that leverages the full potential of deep learning. Our extensive evaluations across diverse datasets demonstrate its superior performance and underscore its potential to enhance our understanding of aging and its biological underpinnings, paving the way for improved diagnostic and therapeutic strategies in age-related diseases.

## Materials and Methods

### Dataset

For this study, we utilized 12 publicly available GEO datasets (Edgar, Domrachev, and Lash 2002) from the Biolearn library (Ying et al. 2023), encompassing a comprehensive age range from newborns to centenarians. The datasets provide rich metadata, including age and sex, allowing for a nuanced analysis of methylation patterns across different demographics. We partitioned these datasets into three groups: 90% of those were kept for training and validation while the remaining samples served as held-out test datasets to evaluate generalizability.

The age distribution across these datasets is depicted in the accompanying Figure 1a, highlighting the mean and standard deviation of ages, which span from 0 to over 100 years. Notably, of the 12 datasets, 8 contain gender information, with a demographic composition of 69% male and 31% female samples, as illustrated in Figure 1b. This gender representation ensures that our findings are robust across both male and female cohorts. Most samples originate from human whole blood tissue, which is commonly used in epigenetic studies due to its accessibility and the abundance of methylation data it offers. This tissue type enhances the relevance and applicability of our study to general and clinical research.

**Figure 1:**
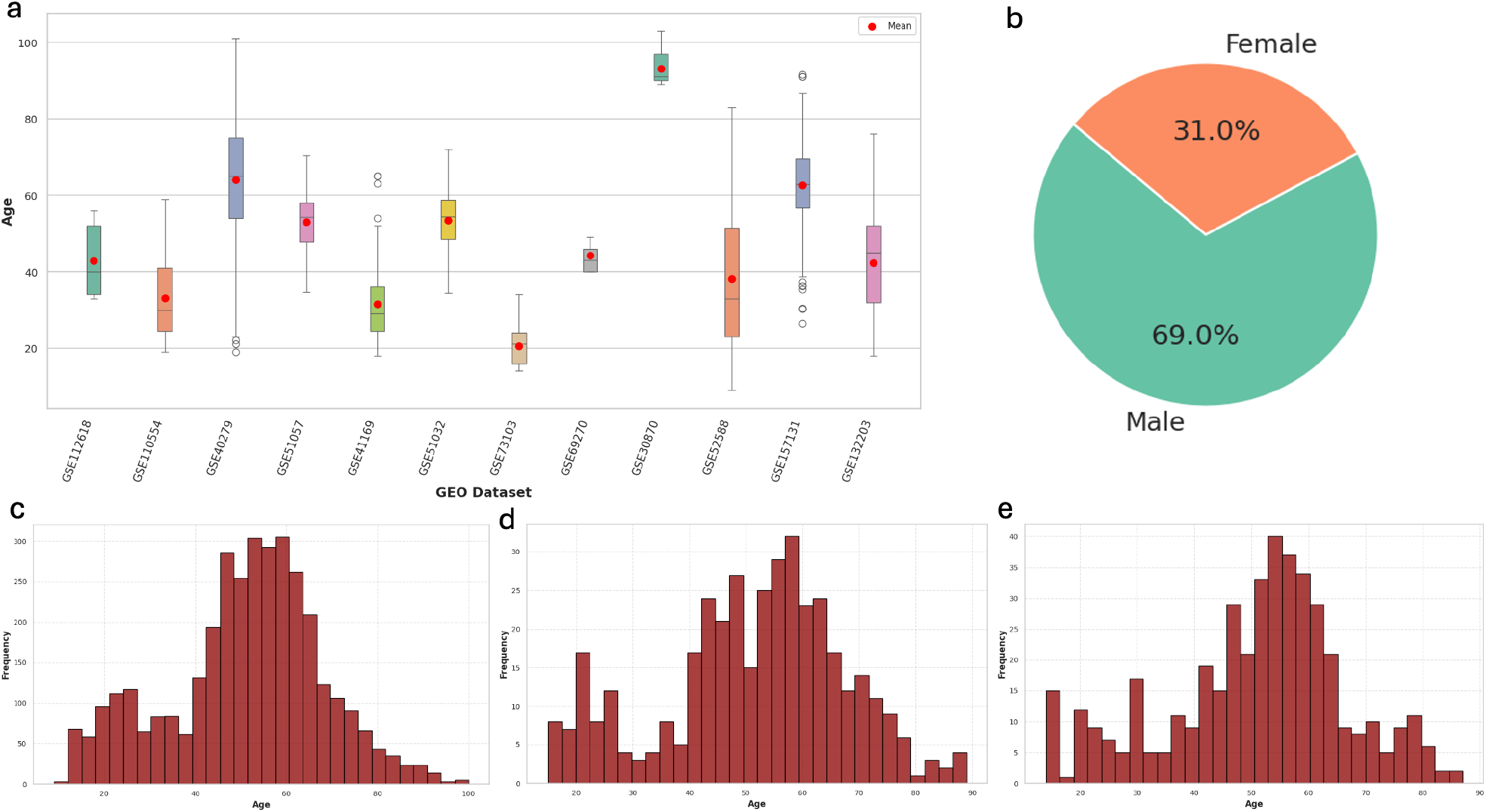
Analysis of GEO Datasets for Age Prediction. a) Age Distribution: A bar plot illustrating the age range from 0 to over 100 years across 12 GEO datasets, with annotations for mean and standard deviation to highlight the diversity in age representation. b) Gender Proportion: Pie chart showing the gender ratio of the sampled populations, restricted to datasets with available sex information. c-e) Data Partitioning: Histograms depicting the age distribution within the training, validation, and test sets, demonstrating consistent distribution patterns across different dataset partitions.

### Data Preprocessing

The preprocessing of our dataset was meticulously uniform across the training, validation, and testing sets to maintain consistency. For the independent test set, the models were assessed directly without further preprocessing to evaluate their performance on unmanipulated data. Initially, we filtered out samples that exhibited more than 50% missing values in their methylation profiles, ensuring the integrity of our dataset. Additionally, any samples lacking accurate age information or presenting with NaN values were excluded. To minimize the influence of extreme age outliers, which could skew the model’s learning process, we removed samples with ages exceeding 100 years, as these were not sufficiently numerous to contribute effectively to model training.

The resulting dataset comprised 4,351 samples, each characterized by 20,937 CpG sites. We then normalized the methylation beta values to a 0 - 1 range using the MinMaxScaler from the scikit-learn library (Pedregosa et al. 2011), ensuring that all values were appropriately scaled for effective model input. Lastly, missing values within the methylation data were imputed using the mean methylation level of each CpG site across all samples, facilitating a consistent dataset for subsequent analyses. This rigorous preprocessing pipeline was essential for preparing the data for accurate and unbiased epigenetic age prediction.

### Age associated Feature Selection and data partitioning

In our study, we implemented a novel Dual-Correlation feature selection technique that significantly enhanced the predictive accuracy of our model. This method integrates both linear and non-linear correlation analyses—specifically Spearman’s rank correlation for detecting monotonic relationships and Pearson’s correlation for linear relationships. By intersecting the significant CpG sites identified by both correlation methods, we were able to capture a comprehensive set of biomarkers reflective of epigenetic aging. This approach not only solidifies the robustness of our feature selection but also provides deeper insights into the complex nature of epigenetic modifications with age.

For correlation thresholds set at T=0.4,0.45,0.5,0.55, and 0.60, we identified 407, 184, 57, 16, and 5 CpG sites, respectively (Selection process is shown in Figure 3a for T=0.45). These varying thresholds allowed us to assess the impact of feature granularity on model performance, ultimately aiding in the optimal selection of predictive biomarkers.

Regarding data partitioning, we initially segregated 10% of the dataset to form an independent test set for final evaluation. The remainder was then divided, reserving 10% for validation purposes. This partitioning strategy resulted in 3,523 training samples, 392 validation samples, and 436 test samples, ensuring that our model was both trained and validated on diverse subsets of the data, promoting generalizability and robust performance across unseen datasets. The age distribution for train, validation and test sets were shown in Figure 1c, d, e respectively.

### Model Architecture

Temporal Convolutional Networks (TCNs) excel in sequence modeling tasks due to their hierarchical architecture, which adeptly manages long-range dependencies. Our implementation incorporates key architectural features that enhance the network’s efficiency and accuracy in age prediction from DNA methylation data shown in Figure 2.

**Figure 2:**
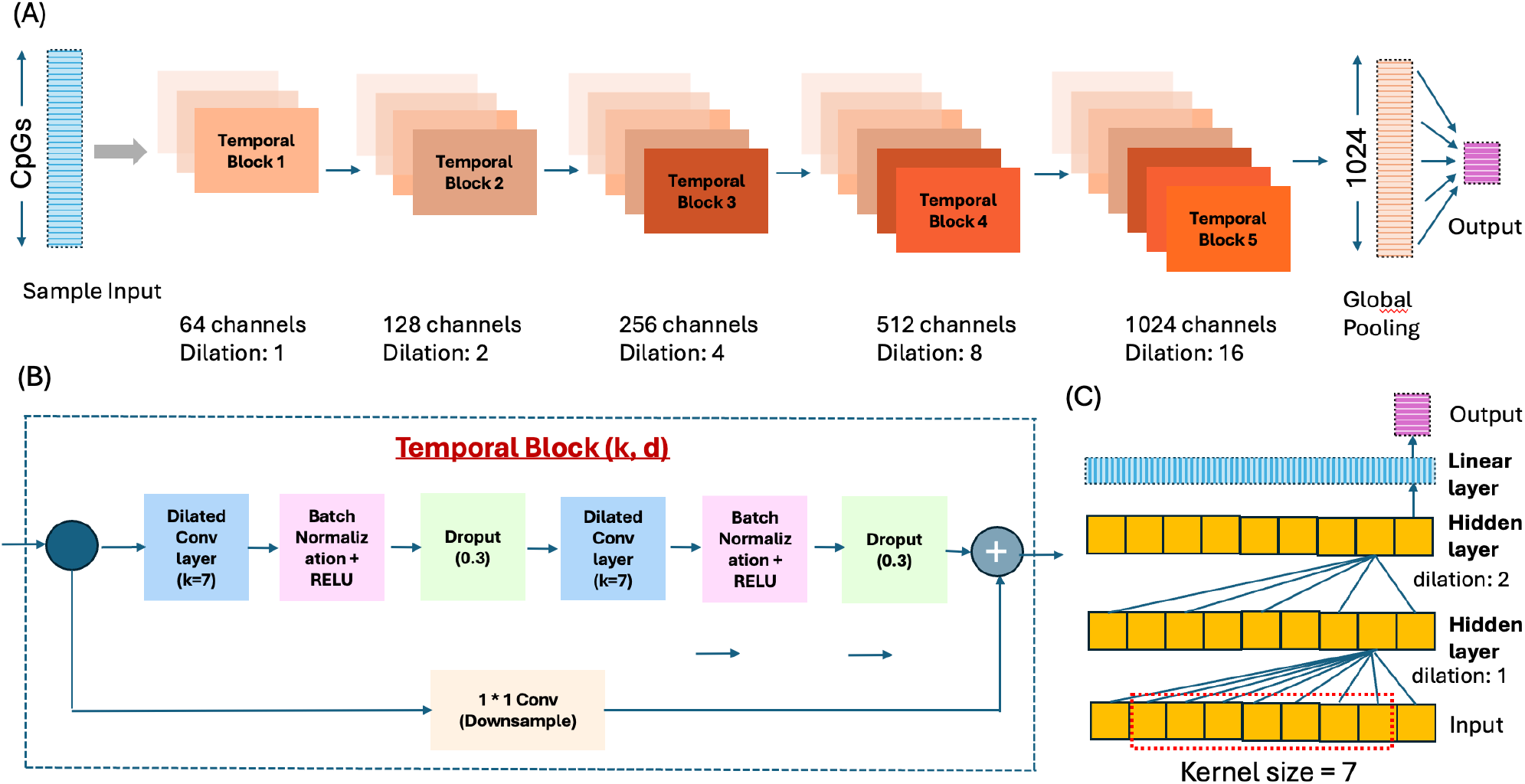
DeepAge Model Architecture. **(A)Input and TCN Overview:** Showcases an input sample with CpG features processed through a TCN with five layers. Each layer doubles the channel count from 64, with exponentially increasing dilations and adaptive padding, culminating in a global max pooling layer that reduces all features to a 1024-dimensional embedding for the output prediction. **(B) Temporal Block Structure:** Details the configuration of a temporal block, including dual convolutional layers followed by batch normalization, ReLU, and dropout, complemented by a skip connection for enhanced gradient flow. **(C) Dilation Mechanics:** Highlights the role of increasing dilation in capturing distant CpG interactions, essential for understanding broader epigenetic patterns.

### Temporal Block

The fundamental unit of our TCN, the TemporalBlock, consists of a series of convolutional layers coupled with ReLU activation (Agarap 2018) and dropout for regularization. Each block features a residual connection, streamlining the training of deep networks by preserving gradient flow and mitigating the vanishing gradient problem. This design ensures robust feature extraction across different layers, facilitating the capture of complex patterns in methylation profiles. Each *TemporalBlock* in our TCN processes the input through two main convolutional layers, each followed by a ReLU activation and dropout, and includes a residual connection:

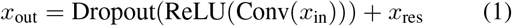

where *x*_res_ is the residual connection that may involve a transformation if the dimensions do not match:

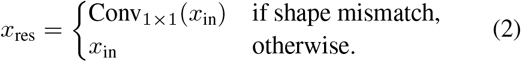

### Dilated Convolutions

To expand the model’s receptive field without a proportional increase in parameters, we employ dilated convolutions. This approach allows the network to integrate information over larger expanses of the input sequence, capturing the distant relationships between CpG sites that are crucial for accurate age estimation. The dilation factor increases exponentially with each subsequent layer, enhancing the model’s ability to assimilate broader contextual information from the methylation data:

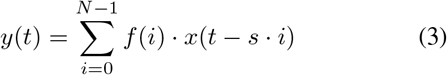

where *s* is the dilation factor, *N* is the filter size, *f* is the filter, and *x* is the input.

### Final TCN Prediction

The overarching model structure, TCNModel, stacks multiple TemporalBlock layers, each refining the feature representations extracted from the data. The architecture concludes with a linear layer that transforms the high-level features into a final age prediction. This layer acts as a regression output, providing a quantitative estimate of biological age:

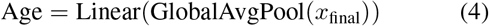

We integrate batch normalization to stabilize the learning process, leading to faster convergence and enhanced training dynamics. Dropout is strategically placed within the TemporalBlocks to prevent overfitting, ensuring that our model generalizes well to new, unseen data. The configuration of these components within the TCN framework allows for a powerful, yet efficient, approach to modeling the intricate relationships inherent in epigenetic data, ultimately leading to more precise age predictions based on DNA methylation patterns.

### Experimental Setup and Model Training

In this study, we developed a temporal convolutional network (TCN) with five layers to predict epigenetic age. Our model architecture included five residual blocks, each comprising two convolutional layers followed by batch normalization and ReLU activation functions. To mitigate overfitting, we introduced a dropout rate of 30% after each convolutional layer. The network’s architecture is designed such that each layer maintains the sequence length of the input (e.g., 57 CpGs), using padding calculated to accommodate the dilation effects. This design allows the number of channels to increase with each layer, enhancing the network’s ability to learn increasingly complex features at deeper levels.

The final layer of the model processes the output with an embedding size that expands to 1024 channels, while preserving the original sequence length. This setup enables the model to represent more complex features without altering the temporal resolution of the input. Following the last temporal block, we applied global average pooling across the sequence dimension, condensing the temporal information into a single vector per feature channel. Consequently, each of the 1024 channels represents the average feature value across all time steps. A linear layer subsequently reduces this 1024-dimensional vector to a single predictive output, focusing on the most critical features for age prediction. This method ensures that the model prioritizes the most significant features extracted throughout the sequence, thereby enhancing its robustness and adaptability to various input lengths.

For training, we employed a batch size of 32, utilized the Adam optimizer with a learning rate of 0.001, and adopted Mean Squared Error as the loss function. The training process was designed to run for up to 200 epochs, with a patience parameter of 5 set for early stopping based on validation loss. The training loss curve is shown in Figure 3b along with validation loss and the training performance is shown in Table 1 along with the validation and test results. This strategy was implemented to prevent overfitting and to halt training when the model ceased to show improvements on the validation dataset.

**Figure 3:**
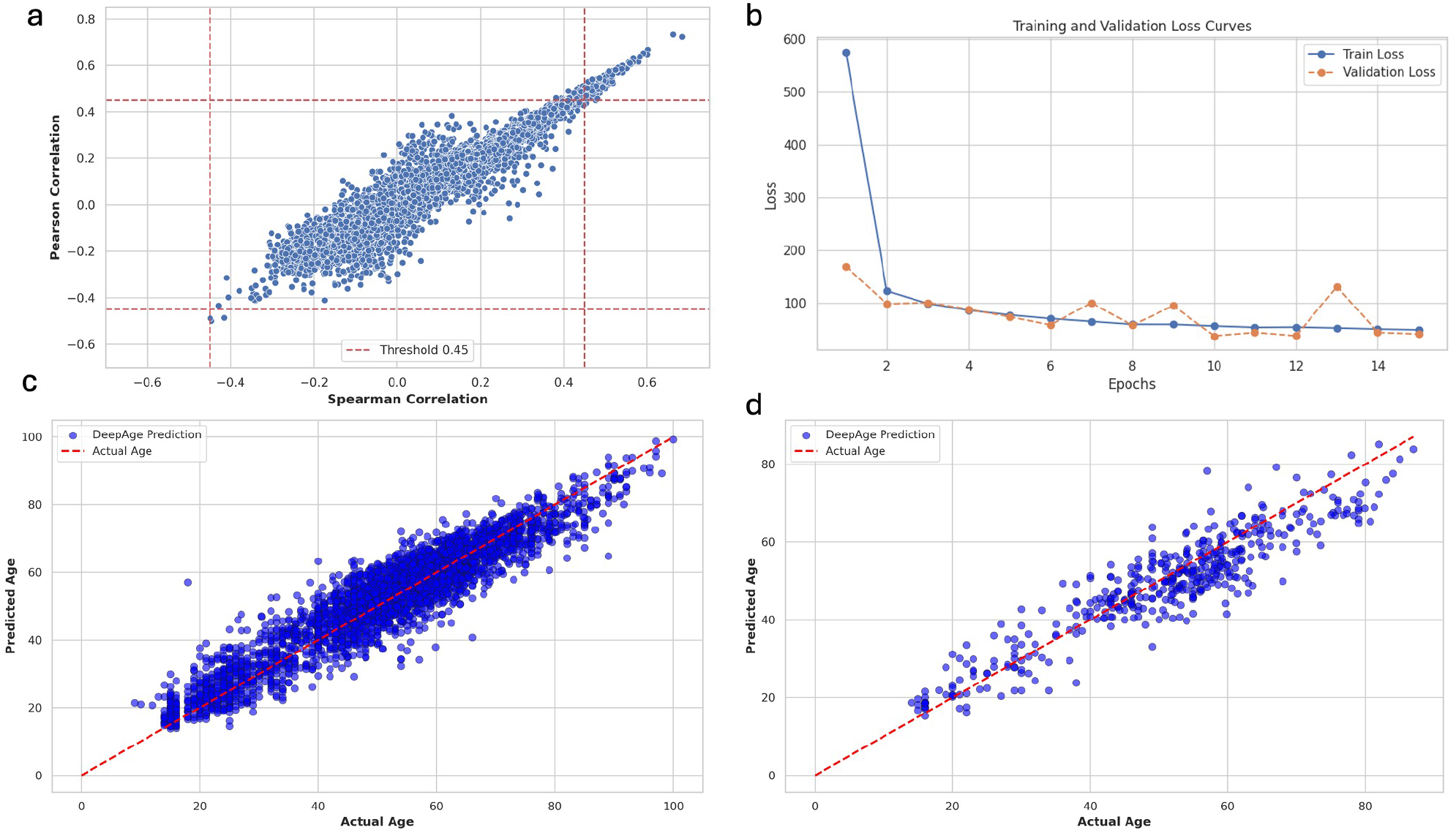
Feature Selection and Model Training. **a) Dual Correlation Method:** Depicts feature selection using a 0.45 correlation threshold. **b) Loss Curves:** Shows training and validation loss curves. **c-d) Age Predictions:** Displays predicted vs. actual age plots for both training and test sets.

**Table 1:** DeepAge performance on train, validation, and test data (using 184 CpGs)

| Metrics | Train | Val | Test |
| --- | --- | --- | --- |
| Mean Absolute Error (MAE) | 4.18 | 4.92 | 4.88 |
| R-Squared ($R^2$) | 0.89 | 0.85 | 0.84 |
| Mean Absolute Deviation (MAD) | 13.19 | 12.98 | 12.23 |
| Root Mean Squared Error (RMSE) | 5.34 | 6.4 | 6.21 |
| Median Absolute Error (MedAE) | 3.52 | 3.86 | 3.98 |

### Evaluation

To effectively measure different model’s age prediction capabilities from DNA methylation data, we employed a robust and comprehensive set of regression metrics. These include:

- Mean Absolute Error (MAE)

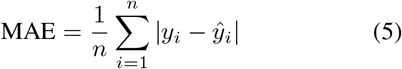
- Root Mean Squared Error (RMSE)

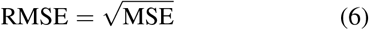
- Median Absolute Error (MedAE)

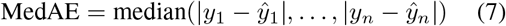
- R-Squared (*R*^2^)

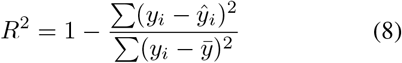
- Mean Absolute Deviation (MAD)

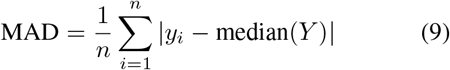

These metrics facilitate a nuanced assessment of prediction accuracy. MAE and RMSE directly reflect average errors in age estimates, with RMSE giving added weight to larger discrepancies. R-squared offers insight into how much of the age-related variance our model captures compared to the baseline model. Collectively, these metrics underscore our model’s ability to generalize well across diverse methylation profiles, substantiating its efficacy in biological age estimation.

## Results

### Comparison of different machine learning and deep learning based methods on age prediction

In our study, we evaluated the performance of various age prediction methodologies, comparing traditional machine learning techniques, ensemble methods, and advanced deep learning architectures against our DeepAge model shown in Table 2. These included regression approaches, gradient boosting, stacking, deep neural networks such as CNN (O’shea and Nash 2015), LSTM (Graves and Graves 2012), and CNN combined with attention mechanisms (Vaswani et al. 2017). For regression based methods and some other models, we utilized the CpG coefficients from the Biolearn library for this comparative analysis, selecting common CpGs between the provided lists and our dataset to facilitate age prediction.

**Table 2:** Performance comparison of different state-of-the-art models on the test dataset.

| Model | No. of CpGs used | MAE | R<sup>2</sup> | RMSE | MedAE |
| --- | --- | --- | --- | --- | --- |
| Hannum | 6 | 34.16 | -4.63 | 37.31 | 32.07 |
| Horvath1 | 297 | 40.56 | -6.61 | 43.34 | 38.55 |
| Horvath2 | 100 | 39.58 | -6.34 | 42.58 | 37.35 |
| PhenoAge | 468 | 40.04 | -5.77 | 40.87 | 39.57 |
| Lin | 89 | 45.69 | -7.77 | 46.55 | 46.63 |
| DunedinPACE | 4 | 40.38 | -5.76 | 43.33 | 38.13 |
| YingAdaptAge | 53 | 58.83 | -13.93 | 60.74 | 56.71 |
| YingDamAge | 50 | 28.31 | -3.19 | 32.18 | 25.58 |
| XGBoost | 57 | 23.77 | -2.22 | 28.21 | 20.42 |
| Random Forest | 57 | 11.57 | 0.12 | 14.77 | 9.03 |
| CNN-Attention | 57 | 11.07 | 0.18 | 14.24 | 8.48 |
| CNN (3 layers) | 57 | 9.26 | 0.49 | 11.25 | 8.43 |
| LSTM (2 layers) | 57 | 7.14 | 0.68 | 8.95 | 6.05 |
| **DeepAge** | **57** | **5.55** | **0.79** | **7.16** | **4.27** |

Notably, we adapted other machine learning and deep learning models from scratch using training data and assessed them on separate test datasets. Our evaluation focused on the efficacy of using 57 CpGs out of a broader set, where DeepAge demonstrated superior performance across all metrics. We also showed the final predicted result by DeepAge vs actual age comparison using 184 CpGs in Figure 3c, 3d for train and test set respectively. This outcome underscores the advantage of treating methylation data as sequential patterns, a strategy that significantly boosts predictive accuracy and is well-illustrated by the success of DeepAge and LSTM-based models. The use of Temporal Convolutional Networks (TCN) in the DeepAge model incorporates crucial architectural elements like residual blocks and dilated convolutions that significantly enhance age prediction accuracy. Residual blocks helped in maintaining the strength of the gradient during backpropagation, allowing for deeper networks without performance degradation. Meanwhile, dilated convolutions expand the receptive field, enabling the model to capture interactions between CpGs that are distant from each other. This capacity to integrate long-range CpG interactions is instrumental in detecting complex patterns in methylation data that are predictive of biological age.

Conversely, convolutional approaches showed moderate effectiveness, suggesting that capturing long-range sequential interactions among CpGs can enhance prediction accuracy. Traditional machine learning methods like random forest and XGBoost underperformed, likely due to their inability to account for complex interactions between CpGs. Additionally, other epigenetic clocks derived from the Biolearn coefficient files exhibited poorer performance, potentially due to the less relevance of their selected CpGs compared to the meticulously curated ones used in our model. This comprehensive analysis highlights the robustness and effectiveness of DeepAge in leveraging epigenetic sequencing for age prediction, setting a new benchmark for accuracy in the field.

### Effect of number of CpG sites and their correlation on age prediction

The performance of age prediction models is significantly influenced by the number of methylation sites, or CpGs, used as features. Including more CpG sites broadens genomic coverage, capturing a diverse array of methylation patterns linked to aging. However, this increased dimensionality can complicate the model, potentially leading to overfitting, where performance is high on training data but poor on unseen data. Furthermore, a larger feature set complicates the interpretation of model predictions, as discerning the most influential CpG sites becomes challenging, complicating biological interpretations and validations.

Not all CpG sites contribute equally to age prediction; some are highly predictive due to their direct biological roles in aging, while others show minimal age-related variations. Thus, employing effective feature selection techniques is crucial to pinpoint the most informative CpGs, enhancing both model performance and computational efficiency.

To determine the optimal number of CpG sites among our samples, we evaluated five different CpG sets selected using Dual-Correlation feature selection at varying thresholds (0.45, 0.50, 0.55, 0.60) shown in Table 3. We found that models with a moderate number of highly associated CpGs perform well, but performance declines beyond a certain point due to overfitting. For example, using 184 CpGs yielded better results than using 407 CpGs, indicating the onset of overfitting. In our analysis, a set of 57 CpGs emerged as a balanced choice, providing robust performance while minimizing complexity.

**Table 3:** Effect of Number of CpGs on Age Prediction.

| Metric | Number of CpGs |  |  |  |  |
| --- | --- | --- | --- | --- | --- |
|  | T=0.40 (407) | T=0.45 (184) | T=0.50 (57) | T=0.55 (16) | T=0.60 (5) |
| MAE | 5.2 | **4.88** | 5.55 | 6.72 | 7.69 |
| R<sup>2</sup> | 0.82 | **0.84** | 0.79 | 0.69 | 0.60 |
| RMSE | 6.62 | **6.21** | 7.16 | 8.62 | 9.88 |
| MedAE | 4.32 | **3.98** | 4.27 | 5.24 | 5.99 |

### Visualization of CpG site methylation levels patterns across samples

In our study, we meticulously analyzed the methylation patterns across various CpG sites to gain insights into their implications for biological age estimation. The heatmap provided visually demonstrates the variation in methylation levels across different samples. Each row represents a sample, and each column corresponds to a specific CpG site. The color gradient, ranging from light yellow (low methylation) to deep red (high methylation), facilitates the identification of CpG sites with pronounced methylation changes. From this visualization, we can discern clear patterns of methylation across specific sites, highlighting regions with potential biological significance in aging processes. Sites with consistently higher or lower methylation across samples could indicate key regulatory regions impacting gene expression tied to aging. This methodical mapping enables us to target these significant CpG sites for deeper analysis, potentially guiding further experimental investigations. Moreover, the heatmap assists in identifying outliers and trends that may not be evident through numerical data alone. In Figure 4a, b we showed the methylation levels for each samples on test and train set respectively across all samples. For example, cg14166009, cg00059225 is highly methylated for bothe train and test samples. By correlating these patterns with age groups and other phenotypic data, we can better understand the role of epigenetic modification in aging and develop more accurate predictive models for biological age. This approach not only enhances the precision of age estimation models but also enriches our understanding of the epigenetic mechanisms that underlie age-related changes.

**Figure 4:**
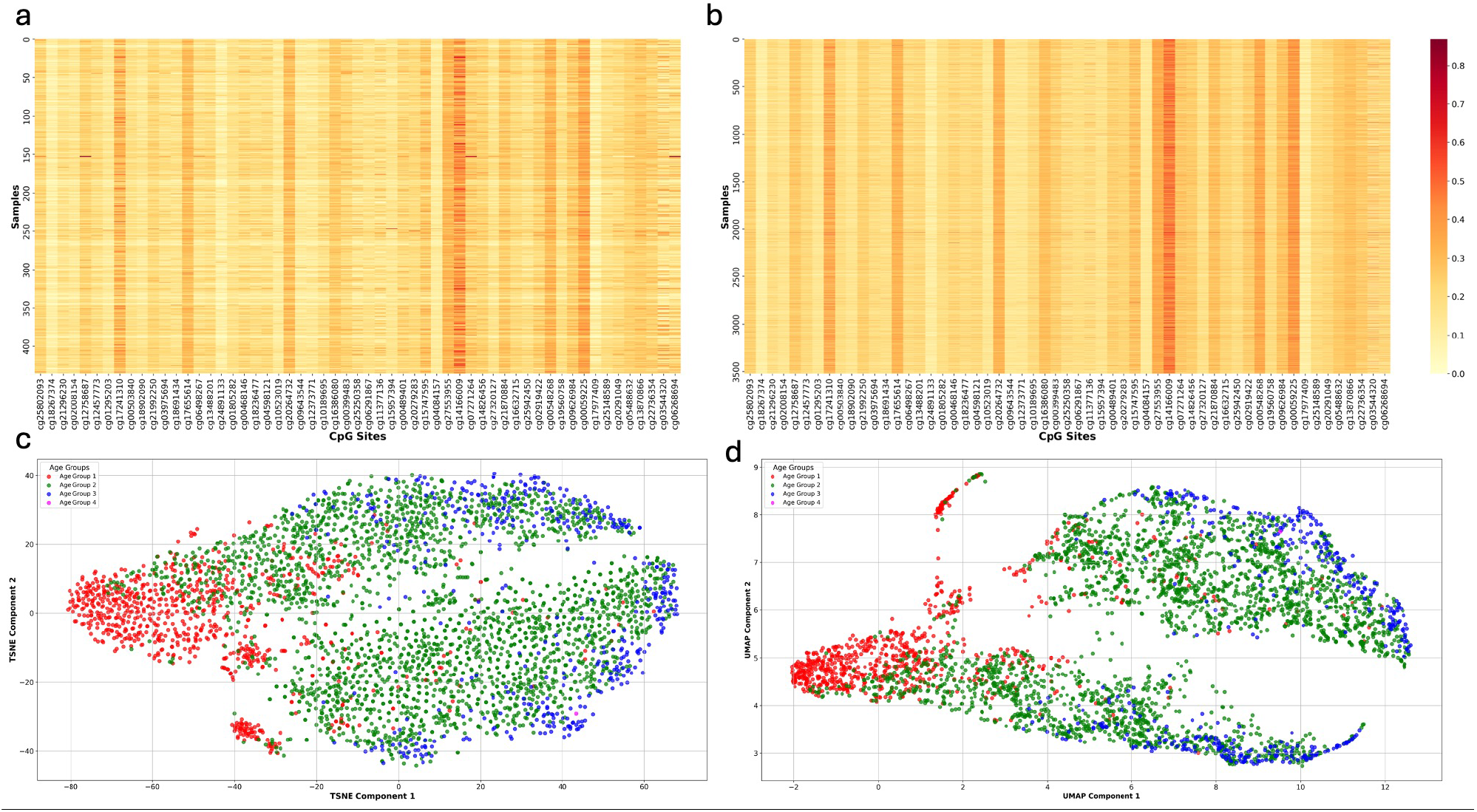
Figure 3: Methylation Data Visualization. a-b) Heatmaps of CpG Sites. Displays methylation levels across samples for test and training sets respectively. **c-d) Dimensionality Reduction:** Illustrates t-SNE and UMAP visualizations of methylation data, categorized into four age groups (0-25, 25-50, 50-75, 75-100) showing distinct patterns for age groups.

### Ablation study on model parameters of DeepAge (57 CpGs)

To optimize our model, DeepAge, for age prediction, we conducted a series of ablation experiments. These experiments were designed to identify the most effective architectural features by varying the number of layers, kernel sizes, and regularization techniques such as batch normalization and dropout. Table 4 below summarizes our findings using the configuration of the 57 CpG model on several key evaluation metrics. This configuration not only highlights the effectiveness of our architectural choices but also underscores the capability of DeepAge to provide reliable and accurate age predictions based on DNA methylation profiles.

**Table 4:** Ablation study on model parameters of DeepAge (57 CpGs)

| Model Parameters | MAE | R<sup>2</sup> | RMSE | MedAE |
| --- | --- | --- | --- | --- |
| TCN k=3, 3 layers, drop=0.2, num_channels = [64, 128, 256] | 9.24 | 0.47 | 11.41 | 7.93 |
| TCN k=7, 3 layers, BN, drop=0.3, num_channels = [64, 128, 256] | 6.47 | 0.73 | 8.1 | 5.45 |
| **TCN k=7, 5 layers, BN, drop=0.3, num_channels = [64, 128, 256, 512, 1024]** | **5.55** | **0.79** | **7.16** | **4.27** |
| TCN k=7, 5 layers, BN, drop=0.3, num_channels = [64, 128, 256, 512, 512, 1024, 1024, 2048] | 12.8 | -0.03 | 15.93 | 10.17 |

Our experiments indicated that increasing the depth of the network through additional layers of residual blocks significantly improved performance up to a certain point. We found that a five-layer model was particularly effective in capturing the complex interactions among methylation sites, providing a balance between model complexity and performance.

We also experimented with various kernel sizes and determined that a kernel size of 7 was optimal. This size allows the model to effectively capture information from neighboring CpG sites, which is crucial to understanding patterns associated with aging. To enhance the model’s generalizability and robustness, we incorporated batch normalization and a dropout rate of 0.3. These techniques help in mitigating the overfitting problem and ensure that the model performs consistently well across different datasets.

### Analysis of variation in DNA methylation and age patterns across different age groups through dimensionality reduction Techniques

In our study, we employed advanced dimensionality reduction techniques, specifically t-SNE (Van der Maaten and Hinton 2008) and UMAP (McInnes, Healy, and Melville 2018), to explore the underlying patterns in DNA methylation data relative to epigenetic age. Our analysis distinctly categorized samples into four age groups, ranging from 0 to 100 years. The visualization results, as illustrated in the accompanying figure, underscore significant differences in methylation patterns across these age groups. The t-SNE visualization in Figure 4c shows a diffuse clustering of samples where some age-based grouping is visible but overlaps significantly between groups. The spread suggests that, while there is some age-related structure in the methylation data, the distinctions between age groups are not sharply defined in this two-dimensional embedding. The UMAP technique, in particular, demonstrated superior capability in delineating age-related differences, showcasing well-detailed clusters that correspond to the predefined age categories shown in Figure 4d. This clear segregation indicates that UMAP is adept at preserving the global structure of the data, providing insightful visualizations that highlight the biological significance of DNA methylation with age. These findings suggest that methylation patterns are not only age-dependent but also can be effectively visualized to reflect age-related biological changes, reinforcing the potential of methylation as a biomarker for biological aging. The improved clustering seen in UMAP highlights its utility in epigenetic studies where understanding the nuanced effects of age on methylation is crucial.

## Discussions

In this study, we developed DeepAge, a novel deep learning framework for estimating epigenetic age using DNA methylation data. By employing Temporal Convolutional Networks (TCNs), our model successfully captures long-range dependencies and interactions between CpG sites, significantly improving upon the performance of traditional epigenetic clocks. DeepAge’s architecture utilizes dilated convolutions to increase the receptive field without a drastic increase in parameters, enhanced by residual connections and dropout for robust learning even in the presence of limited data.

However, our study does have limitations that open avenues for future research. We restricted our analysis to 12 datasets from the BioLearn library, focusing on human blood samples which, while diverse, represent a constrained view of possible epigenetic aging patterns. The data, covering a broad range of ages, sexes, and ethnicities, provides a solid basis for age prediction but is still limited in scale and variety. Expanding our dataset to include more samples, especially from underrepresented groups or additional datasets outside of BioLearn, could enhance the model’s accuracy and generalizability. Expanding our model to include data from multiple tissues could enhance its versatility and relevance across diverse biomedical queries. This broader approach may improve prediction accuracy and provide detailed insights into the distinct aging patterns of various tissues, potentially informing more targeted age-related disease research and interventions.

Despite these limitations, DeepAge represents a significant advancement in the field of epigenetic age estimation. It showcases the potential of deep learning to elucidate complex biological relationships and highlights the importance of innovative computational methods in the ongoing exploration of aging and regenerative medicine. Future work will focus on expanding the dataset and refining the model architecture to further enhance its performance and applicability.

## Code and Data Availability

Code will be released during/after the review period. We have used publicly available data from NCBI GEO database (https://www.ncbi.nlm.nih.gov/gds) from biolearn library (https://bio-learn.github.io/data.html)

